# Suppressors of YpsA-mediated cell division inhibition in *Bacillus subtilis*

**DOI:** 10.1101/2020.02.12.946632

**Authors:** Robert S. Brzozowski, Brooke R. Tomlinson, Michael D. Sacco, Judy J. Chen, Anika N. Ali, Yu Chen, Lindsey N. Shaw, Prahathees J. Eswara

**Affiliations:** Department of Cell Biology, Microbiology and Molecular Biology, University of South Florida, Tampa, FL 33620, USA; Department of Molecular Medicine, University of South Florida, Tampa, FL 33612, USA

**Keywords:** FtsZ, GpsB, filamentation, SLOG, YfhS

## Abstract

Although many bacterial cell division factors have been uncovered over the years, evidence from recent studies points to the existence of yet to be discovered factors involved in cell division regulation. Thus, it is important to identify factors and conditions that regulate cell division to obtain a better understanding of this fundamental biological process. We recently reported that in the Gram-positive organisms *Bacillus subtilis* and *Staphylococcus aureus*, increased production of YpsA resulted in cell division inhibition. In this study, we isolated spontaneous suppressor mutations to uncover critical residues of YpsA and the pathways through which YpsA may exert its function. Using this technique, we were able to isolate four unique intragenic suppressor mutations in *ypsA* (E55D, P79L, R111P, G132E) that rendered the mutated YpsA non-toxic upon overproduction. We also isolated an extragenic suppressor mutation in *yfhS*, a gene that encodes a protein of unknown function. Subsequent analysis confirmed that cells lacking *yfhS* were unable to undergo filamentation in response to YpsA overproduction. We also serendipitously discovered that YfhS may play a role in cell size regulation.

**GRAPHICAL ABSTRACT:** 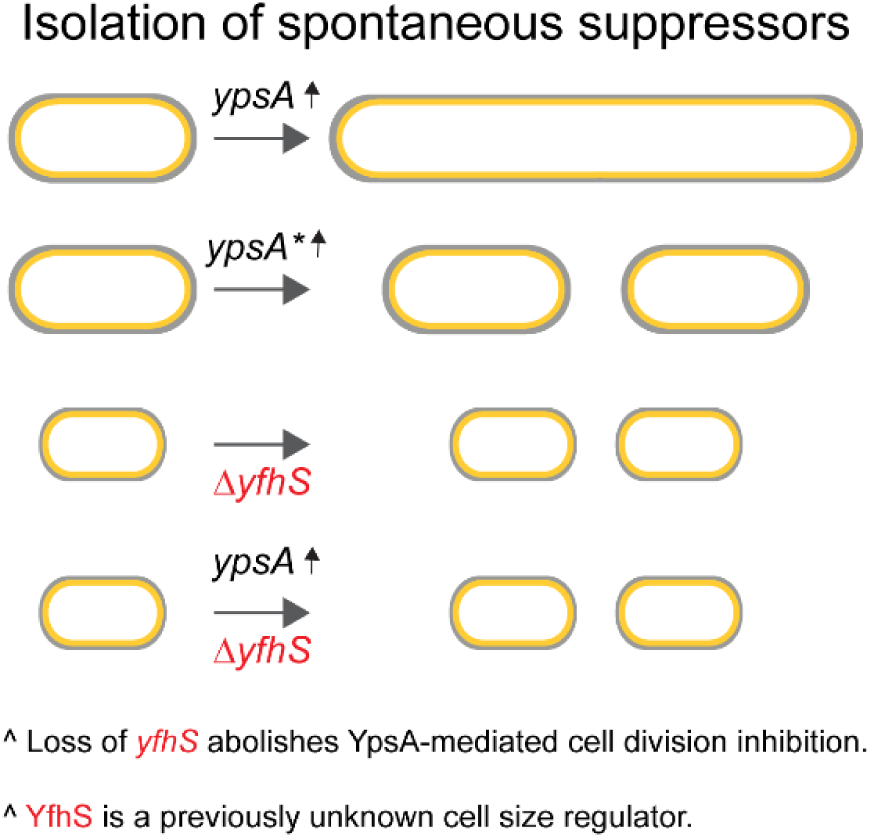

**ABBREVIATED SUMMARY:** In *Bacillus subtilis*, we discovered that increased expression of *ypsA* results in cell division inhibition and impairment of colony formation on solid medium. Colonies that do arise possess compensatory suppressor mutations. Analysis of one such suppressor mutation led us to a protein of unknown function, YfhS, which appears to play a role in regulating cell length and cell width.

## INTRODUCTION

Bacterial cell division is an essential process orchestrated by a multitude of cell division proteins (Haeusser & Margolin, 2016). During growth an essential cell division protein FtsZ, forms a ring-like structure and marks the site of division. There it serves in the recruitment of additional divisome proteins and commences septation (Du & Lutkenhaus, 2017, Errington & Wu, 2017). Although known FtsZ regulatory systems, such as the Min system and nucleoid occlusion, have been well characterized (Eswara & Ramamurthi, 2017), recent studies have determined that correct cell division site selection can occur independent of these mechanisms in both *Bacillus subtilis* and *Escherichia coli* (Rodrigues & Harry, 2012, Bailey *et al.*, 2014). These findings highlight the need to investigate and discover other factors involved in regulating cell division in bacteria. In our lab, we have identified a potential cell division regulator in *B. subtilis* and *Staphylococcus aureus*, YpsA (Brzozowski *et al.*, 2019a).

YpsA is conserved in the Firmicutes phylum of Gram-positive bacteria and appears to be in a syntenous relationship with a known cell division protein, GpsB (Brzozowski *et al.*, 2019a). The crystal structure of *B. subtilis* YpsA was solved by a structural genomics group in 2006 [PDB ID: 2NX2; (Ramagopal *et al.*, 2006)]. Based on the structural features, YpsA was placed as the founding member of the “YpsA proper” subclade within the SLOG (<u>S</u>MF/DprA/*<u>LOG</u>*) protein superfamily (Burroughs *et al.*, 2015), yet the precise function of YpsA remains to be elucidated. The structure of YpsA resembles that of DprA (RMSD with DprA of *Helicobacter pylori* is 2.79 Å, PDB ID: 4LJR), another member of the SLOG superfamily, which is a single-stranded DNA-binding protein involved in DNA recombination (Yadav *et al.*, 2014, Wang *et al.*, 2014). Previously we had found that YpsA provides oxidative stress protection in *B. subtilis* and that overproduction of YpsA results in cell division inhibition, through FtsZ mislocalization, in a growth rate dependent manner (Brzozowski *et al.*, 2019a). Additionally, using site-directed mutagenesis we identified multiple amino acid residues that are potentially important for the structure and/or function of YpsA, including residues located in the conserved substrate binding pocket made up of glycine and glutamate residues predicted by Burroughs *et al*., 2015 (Burroughs *et al.*, 2015). In addition, we have shown that the function of YpsA in cell division is also conserved in the Gram-positive pathogen, *S. aureus* (Brzozowski *et al.*, 2019a).

In this study we utilized a classic spontaneous suppressor isolation technique for further identification of critical amino acid residues that are important for YpsA structure and/or function, and for the elucidation of the molecular mechanism through which YpsA acts. By screening for suppressor mutations of a lethal YpsA overproduction phenotype, we were able to isolate and characterize four unique intragenic suppressor mutations (E55D, P79L, R111P, G132E). Each of these mutations was found to prevent lethality and related cell division inhibition. In addition, we also identified an extragenic suppressor mutation that introduced a premature stop codon in the *yfhS* gene, which codes for a protein of unknown function. Upon subsequent analysis, we verified that in cells lacking *yfhS*, YpsA-dependent cell division inhibition is abolished. Here, we speculate the possible ways by which YfhS and YpsA-mediated cell division phenotypes could be linked. Interestingly, during the course of our experiments, we discovered that YfhS may play a role in cell size regulation, as *yfhS* null cells are significantly smaller in cell width and length when compared to the wild type control.

## RESULTS

### Overexpression of *ypsA* results in a growth defect on solid medium

We have previously shown that overproduction of either YpsA or YpsA-GFP results in severe filamentation in *B. subtilis* [(Brzozowski *et al.*, 2019a); **Fig. 1A**)], a phenotype that is characteristic of cell division inhibition in this organism. To test whether filamentous growth in the presence of inducer results in a distinguishable phenotype on solid medium, we conducted a spot assay. Briefly, serial dilutions of exponentially growing wild type (WT) cells, and cells containing an IPTG-inducible copy of either *ypsA* or *ypsA-gfp* were spotted on solid growth medium with or without inducer. In the absence of inducer all strains of all tested dilutions grew similar to the WT control (**Fig. 1B**; see left panel). In the presence of inducer, we observed a significant growth defect associated with both YpsA and YpsA-GFP overproduction, suggesting that cell division inhibition caused by *ypsA* or *ypsA-gfp* overexpression was lethal (**Fig. 1B**; see right panel). Interestingly, YpsA-GFP overproduction resulted in a more severe growth phenotype compared to untagged YpsA. To test whether this difference is due to increased accumulation of YpsA-GFP in the cells, we tagged both YpsA and YpsA-GFP with a FLAG tag at their C-terminus and conducted an anti-FLAG western blot analysis. Overproduction of YpsA-FLAG and YpsA-GFP-FLAG resulted in filamentation and a growth defect on solid medium that was of similar extent when compared to their non-FLAG tagged counterparts (**Figs. S1A** and **S1B**). The ratio of YpsA-GFP-FLAG and SigA (internal loading control) was similar to that of YpsA-FLAG and SigA (0.68 and 0.80 respectively; **Fig. S1C**), suggesting that the increased lethality of the GFP-tagged version is not due to any changes in accumulation. Next, we utilized the severe lethality elicited by YpsA-GFP overproduction as a tool to isolate spontaneous suppressors.

**Figure 1.**
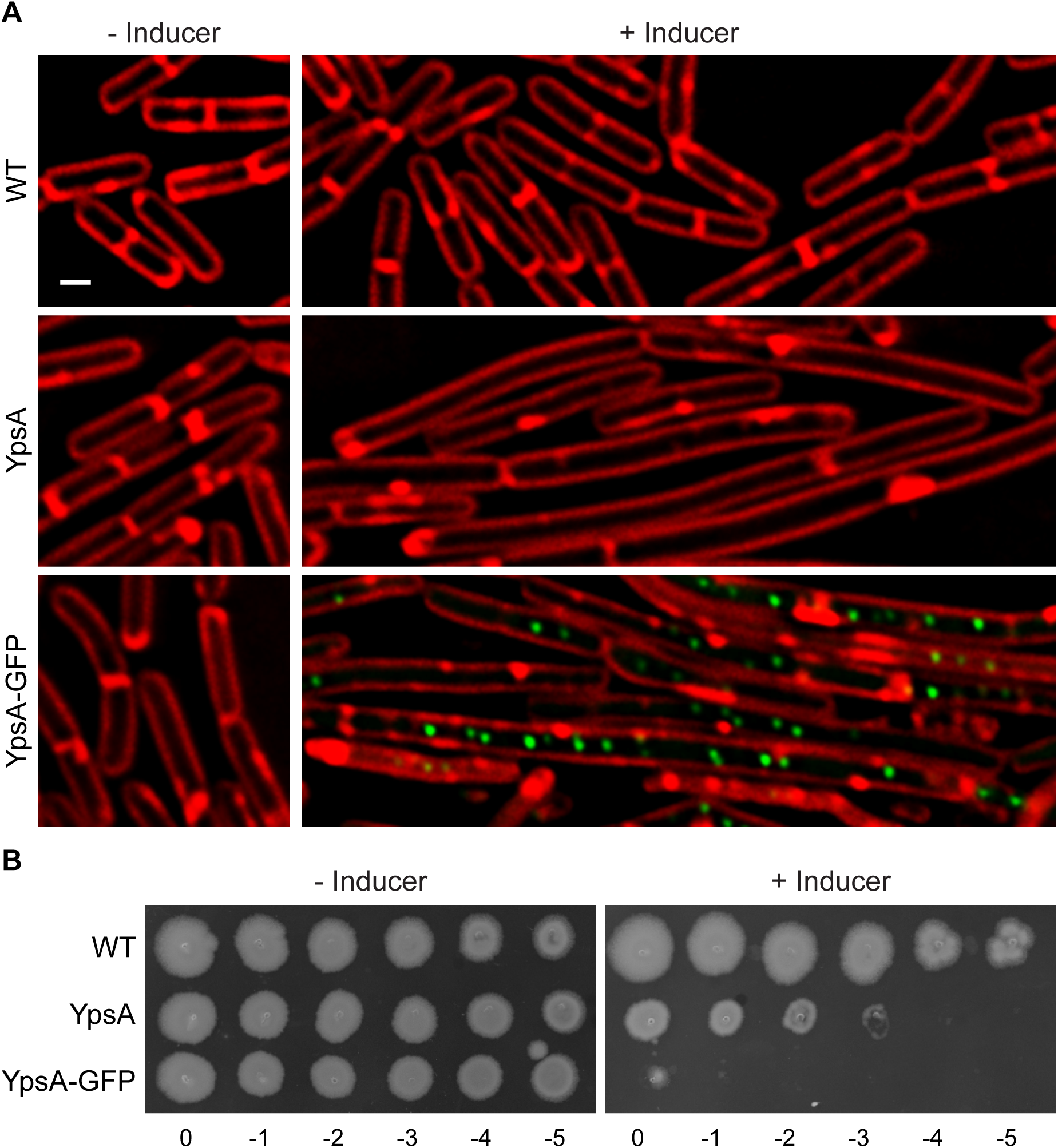
Overproduction of YpsA or YpsA-GFP results in a lethal phenotype. (**A**) Cell morphology of wild type (PY79; WT) cells and cells harboring an IPTG-inducible copy of either *ypsA* (GG82) or *ypsA-gfp* (GG83) grown in the absence of inducer or in the presence of 250 μM IPTG. Fluorescence of red membrane stain (FM4-64; red) and GFP (green) are shown. Scale bar: 1 μm. (**B**) Spot assays of WT cells (PY79) and cells containing an inducible copy of either *ypsA* (GG82) or *ypsA-gfp* (GG83). Serially diluted standardized cultures were spotted on plates containing no inducer (left panel) or 1 mM IPTG (right panel) and grown overnight at 37 °C. Corresponding dilution factors are indicated below.

### Isolation of spontaneous suppressor mutations

The YpsA-GFP overproducing strain was streaked out for single colony isolation on multiple inducer containing plates and the plates were incubated overnight as described in the methods section. Only a few colonies formed per plate, presumably due to spontaneous suppressor mutations, which allow for normal growth despite the presence of inducer. After multiple iterations of suppressor isolation, likely mutations were subsequently determined to be either intragenic (within inducible *ypsA-gfp*, henceforth noted as *ypsA*\**-gfp* for simplicity) or extragenic (elsewhere on the chromosome), and their chromosomes were sequenced to identify the mutations (**Fig. S2**). Using this approach, we were able to isolate four unique intragenic suppressor mutations: G132E, P79L, R111P, and E55D (listed in the order of isolation). Immunoblotting indicated that these mutant versions of YpsA were stably produced (**Fig. 2L**). Next, fluorescence microscopy was used to determine whether these mutations were able to rescue the lethal filamentous phenotype observed when unmutated YpsA-GFP was overproduced. This was carried out on exponentially growing cells of the *ypsA-gfp* overexpression strain and all *ypsA*\**-gfp* intragenic suppressor strains in the absence or presence of inducer. In the absence of inducer all strains exhibited similar cell lengths [YpsA-GFP: 3.23 ± 0.74 μm (**Fig. 2A**); G132E: 3.49 ± 0.93 μm (**Fig. 2C**); P79L: 3.24 ± 0.81 μm (**Fig. 2E**); R111P: 3.16 ± 0.76 μm (**Fig. 2G**); E55D: 3.23 ± 0.83 μm (**Fig. 2I**) | n=100 for all cell length measurements]. Upon the addition of inducer, we found that cells overexpressing *ypsA*\**-gfp* did not exhibit filamentation, unlike the *ypsA-gfp* control [YpsA-GFP: 8.93 ± 5.67 μm (**Fig. 2B**); G132E: 2.99 ± 0.88 μm (**Fig. 2D**); P79L: 3.29 ± 0.81 μm (**Fig. 2F**); R111P: 3.06 ± 0.98 μm (**Fig. 2H**); E55D: 3.10 ± 0.83 μm (**Fig. 2J**)], indicating that the intragenic suppressors were unable to elicit filamentation upon overproduction. We also noted that G132E, P79L, and R111P suppressors displayed impaired foci formation in comparison to the *ypsA-gfp* control (**Figs. 2B, 2D, 2F**, and **2H**). However, foci formation in E55D suppressor was not impaired (**Fig. 2J**).

**Figure 2.**
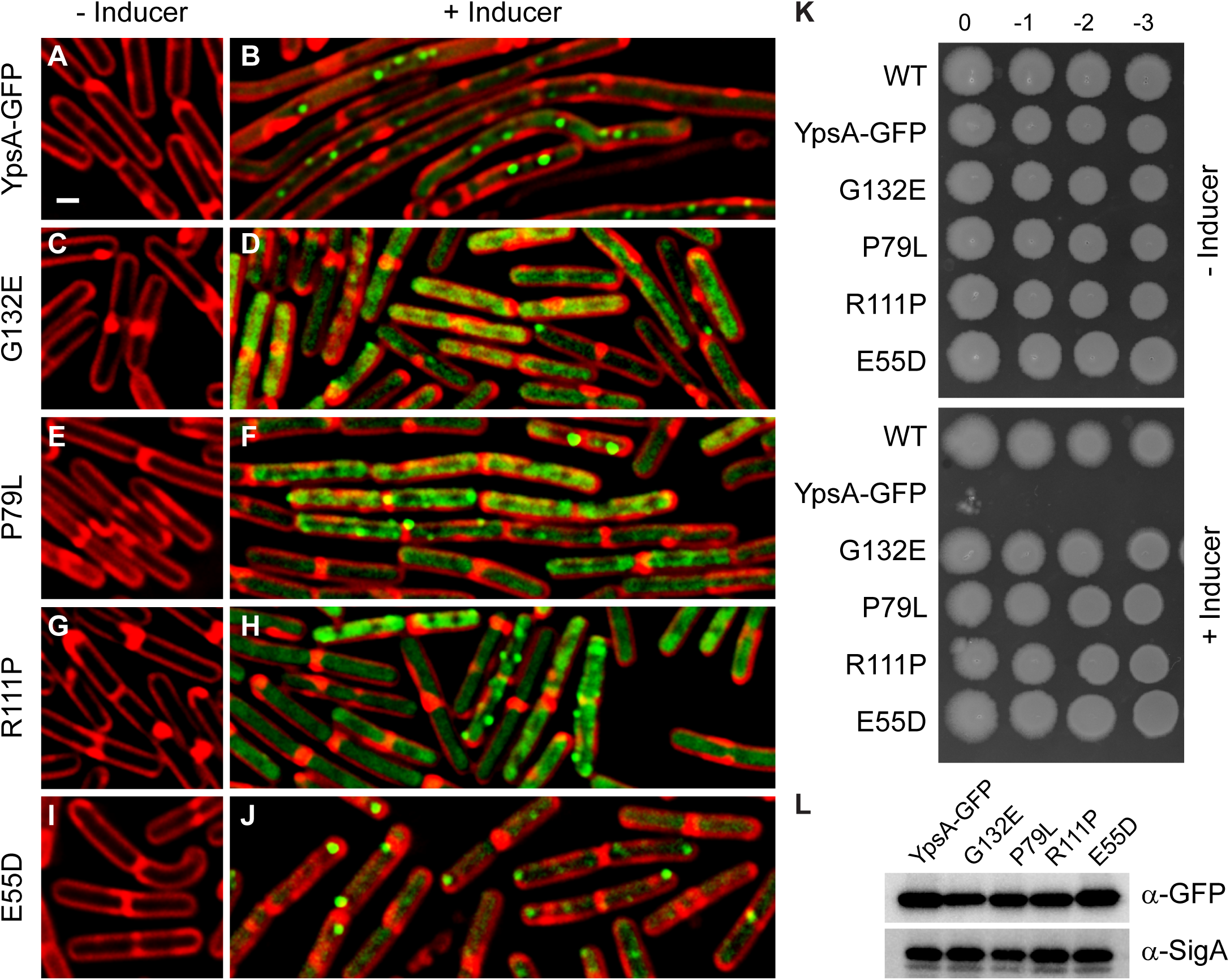
Isolation of spontaneous suppressors. (**A-J**) Fluorescence microscopy comparing cells containing an inducible copy *ypsA*-*gfp* (GG83) and cells containing intragenic mutations (*ypsA*\**-gfp*) isolated during the suppressor screen that resulted in single amino acid changes: G132E (RB300), P79L (RB301), R111P (RB328), and E55D (RB327). Cells were grown in the absence (**A, C, E, G, I**) or in the presence (**B, D, F, H**, of 250 μM IPTG. Fluorescence of FM4-64 (red) and GFP (green) are shown. Scale bar: 1 μm. (**K**) Spot assays of strains harboring an IPTG inducible copy of *ypsA-gfp* (GG83) or *ypsA*\**-gfp* (RB300, RB301, RB328, RB327) grown without inducer (top panel) or with 1 mM IPTG (bottom panel). Corresponding dilution factors are shown on top. (**L**) Stability of YpsA-GFP and YpsA*-GFP variants were confirmed when cells were grown in the presence of inducer. Cell lysates were probed via immunoblotting using anti-GFP and anti-SigA (loading control) antisera.

Each of the intragenic suppressors were subjected to a spot assay to test whether these point mutations were able to grow normally in contrast to the *ypsA-gfp* overexpression strain that displayed a lethal phenotype on solid medium in the presence of inducer. In the absence of inducer all strains grew similar to the WT control (**Fig. 2K**; see top panel). When grown in the presence of inducer, overexpression of *ypsA-gfp* resulted in a severe growth defect (**Fig. 2K**; see bottom panel). However, growth was similar to WT in all intragenic suppressor strains when grown in the presence of inducer (**Fig. 2K**; see bottom panel). Given that these mutants were unable to cause filamentation, it appears that the lethality is directly linked to the ability of YpsA to elicit filamentation. Collectively, these data indicate that the residues E55, P79, R111, and G132 are critical for the function of YpsA, especially in regard to cell division inhibition.

### Structural analysis of the intragenic suppressor mutations

Three of the four YpsA mutants have residues that are buried in the core: E55D, G132E, and P79L (**Fig. 3A**). When large mutations occur in this environment, misfolding and loss of function is often the consequence (Baruah & Biswas, 2014). Amongst these mutants, P79 is significant because it, and its adjacent residue F80, are strictly conserved amongst the YpsA clade of Firmicutes (Burroughs *et al.*, 2015, Brzozowski *et al.*, 2019a). The crystal structure of *B. subtilis* YpsA (PDB ID: 2NX2) reveals P79 and F80 also line the possible DNA binding groove of YpsA (Ramagopal *et al.*, 2006). The positioning of an aromatic side chain here suggests it may facilitate DNA binding by forming stacking interactions with nucleobases (Baker & Grant, 2007). The P79L mutation not only creates severe clashes with surrounding residues, but likely perturbs the positioning of F80, and therefore may directly impair DNA or nucleotide binding (**Fig. 3B**). Likewise, the E55D mutant affects a second, highly conserved segment of the putative DNA binding domain, the GxE motif. In YpsA, E55 is highly coordinated by five potential hydrogen bonds with the side chain and backbone of S49, the T7 sidechain, and the backbone amide of Q51 (**Fig. 3C**). Considering the size and physicochemical properties of their sidechains, one would expect an E→D mutation to have a non-deleterious effect on YpsA. However, in this instance, shortening the sidechain of E55 by a methylene results in the weakening or total loss of these hydrogen bonds and likely destabilizes the possible DNA binding groove. In addition, the aliphatic part of the E55 side chain forms extensive van der Waals interactions with nearby residues such as Q51 and others, and the carboxylate group of an aspartate residue at this position would also clash with these surrounding residues. The third core mutation, G132E, is located at the beginning of a β-strand and is surrounded by multiple bulky and hydrophobic residues including Y164, P162, L1, L4, and F38 (**Fig. 3D**). Conversion from glycine to any other residue besides alanine results in clashes that will affect the secondary structural elements from which the surrounding residues originate. Indeed, every possible G132E rotamer produces significant clashes, with interatomic distances less than 2.2 Å.

**Figure 3.**
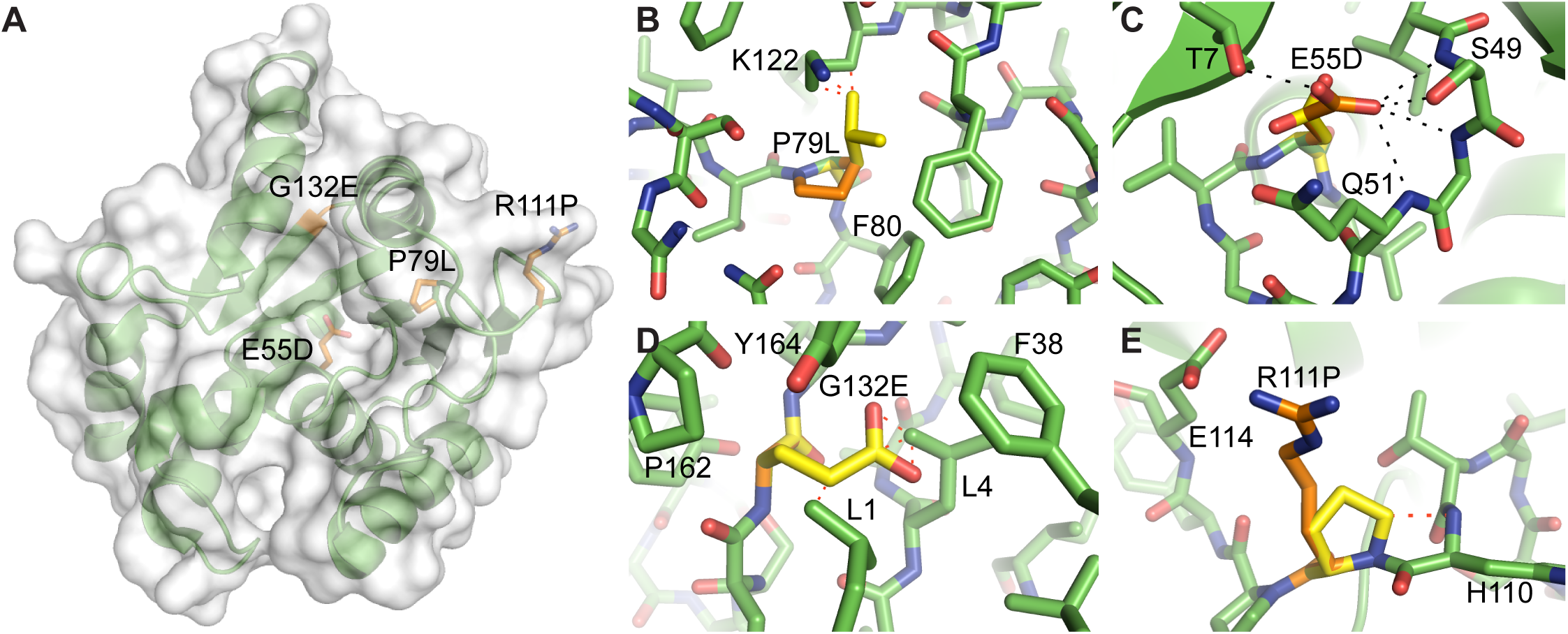
Structural analysis of the intragenic suppressors. (**A**) Crystal structure of *B. subtilis* YpsA (PDB ID: 2NX2) with sites of mutation colored in orange. (**B-E**) Computationally generated mutants are shown in yellow. Hydrogen bonds are shown as black dashes, while steric clashes are represented as red dashes. For clarity, only the most severe clashes are indicated with interatomic distances less than 2.2 Å. (**B**) The P79L mutation generates severe clashes with multiple surrounding residues. (**C**) The E55D mutation in the putative DNA-binding groove results in the potential loss of five hydrogen bonds, destabilizing this region. (**D**) The G132E mutation, similar to P79L mutant, involves a core residue that cannot accommodate any large side chains without severe steric clashes. (**E**) The R111P mutant eliminates a salt bridge with E114 and produces a clash with the adjacent H110 backbone. As a surface residue, it also potentially disrupts intermolecular interactions and signaling.

The R111P mutation is the only intragenic suppressor mutation that involves a solvent exposed residue. Here R111 normally forms a salt bridge with the neighboring E114 (**Fig. 3E**). The conversion from a positively charged side chain to a nonpolar one eliminates this interaction. Furthermore, the cyclic nature of the proline side chain introduces a steric clash with the backbone amide nitrogen of its adjacent residue, H110. The R111P mutation will likely force conformational changes in the protein backbone and cause significant disruptions in intramolecular interactions involving nearby residues. A second scenario that leads to the disruption of YpsA function in this mutant involves the impairment of intermolecular interactions and macromolecular recognition. It is possible that a mutation from R→P prevents interaction with other protein partners of YpsA.

### Isolation and validation of an extragenic suppressor mutation in *yfhS*

Using the same suppressor screening approach (**Fig. S2**), we were able to isolate and validate an extragenic suppressor mutation, which is a duplication of a stretch of 10 nucleotides that introduces a premature stop codon in *yfhS* (**Fig. S3A**). YfhS is a 74 amino acid containing protein of unknown function. *yfhS* is annotated as a sporulation gene upregulated by SigE sigma factor during sporulation (Zhu & Stulke, 2018). However, there is no sporulation defect in a *yfhS* null strain (Yamamoto *et al.*, 1999). *yfhS* may also be regulated by the transcription factor AbbA (Banse *et al.*, 2008).

To test whether disruption of *yfhS* restores normal cell length in cells overexpressing either *ypsA* or *ypsA-gfp*, we generated a strain harboring an inducible copy of either *ypsA* or *ypsA-gfp* in a *yfhS* null background. These strains were then screened with the appropriate controls via a spot assay in order to observe whether or not the *yfhS* deletion was able to restore normal growth on solid medium even when YpsA or YpsA-GFP was overproduced. In the absence of inducer, WT cells and cells containing an inducible copy of either *ypsA* or *ypsA-gfp* grew similarly (**Fig. 4A**). Cells lacking *yfhS* formed small colonies in comparison to WT suggesting an intrinsic growth phenotype associated with the deletion of *yfhS*. Cells harboring an inducible copy of *ypsA* or *ypsA-gfp* in a *yfhS* null background grew similar to the *yfhS* null control strain. When grown in the presence of inducer, as shown in **Fig. 1B**, cells harboring an inducible copy of *ypsA* showed a moderate growth defect while inducible *ypsA-gfp* strain exhibited a severe growth defect (**Fig. 4B**). In the presence of inducer, cells harboring a *yfhS* knockout and cells harboring a *yfhS* knockout with an inducible copy of either *ypsA* or *ypsA-gfp* grew similarly, suggesting that deletion of *yfhS* prevents elicitation of lethal phenotypes displayed by YpsA or YpsA-GFP overproducing cells (**Fig. 4B**). To ensure the phenotype was specific to the disruption of the native copy of *yfhS*, we introduced *yfhS* at an ectopic locus under an inducible promoter. In this complementation strain, the presence of inducer or even leaky expression in the absence of inducer, restored WT-like growth (**Fig. 4A**). Interestingly, the defective growth phenotype of YpsA and YpsA-GFP overproducing cells was also restored in the presence of inducer in the complementation strain (compare **Figs. 4A** and **4B**).

**Figure 4.**
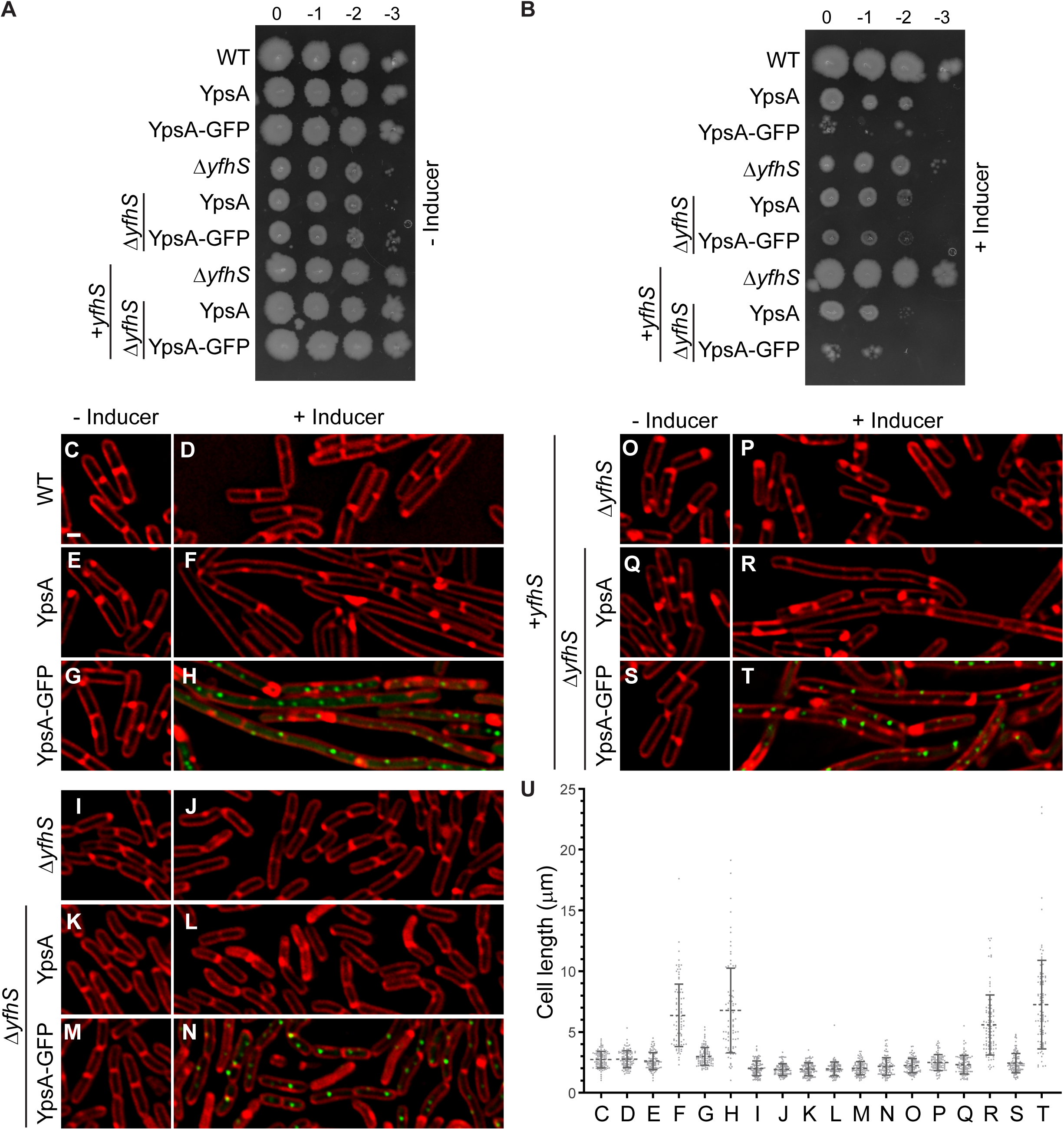
Deletion of *yfhS* rescues YpsA-mediated toxicity and associated filamentation. (**A-B**) Spot assay of WT cells (PY79), Δ*yfhS* cells (RB314), Δ*yfhS* + *yfhS* (RB409) cells, and cells overexpressing either *ypsA* or *ypsA-gfp* in an otherwise wild type background (GG82 and GG83), a Δ*yfhS* background (RB288 and RB289), or in a Δ*yfhS* complementation strain where an intact copy of *yfhS* is engineered to be under the control of an IPTG-inducible promoter at an ectopic locus (RB410 and RB411). Cultures were standardized and serial dilutions were spotted on solid medium without inducer (**A**) or with 1 mM IPTG (**B**). Corresponding dilution factors are indicated on top. (**C-T**) Fluorescence microscopy comparing cell morphologies of WT cells (PY79), Δ*yfhS* cells (RB314), Δ*yfhS* + *yfhS* (RB409) cells, and cells overexpressing either *ypsA* or *ypsA-gfp* in an otherwise wild type background (GG82 and GG83), a Δ*yfhS* background (RB288 and RB289), or in a Δ*yfhS* complementation strain (RB410 and RB411). Cells were imaged in the absence of inducer (**C, E, G, I, K, M, O, Q, S**) or in the presence of 250 μM IPTG (**D, F, H, J, L, N, P, R, T**). Fluorescence signal of FM4-64 membrane dye (red) and GFP (green) are shown. Scale bar: 1 μm. (**U**) Cell lengths of strains shown in panels **C-T** were quantified. The corresponding mean value and standard deviations (n=100) are shown.

Next, we inspected the cell morphology of all strains tested in **Figs. 4A** and **4B** through fluorescence microscopy. Cell division in cells harboring an inducible copy of either *ypsA* or *ypsA-gfp*, but not in WT control, were inhibited upon the addition of inducer (Fig. **4C-H**), as discussed previously [**Fig. 1A**; (Brzozowski *et al.*, 2019a)]. The quantification of cell lengths are shown in **Fig. 4U**. The cell lengths of the WT control strain in the absence and presence of inducer were similar [WT (- inducer): 2.72 ± 0.68 μm (**Fig. 4C**); WT (+ inducer): 2.77 ± 0.69 μm (**Fig. 4D**)]. On the contrary, as expected, cells overproducing YpsA or YpsA-GFP exhibited filamentation in the presence of inducer [YpsA (- inducer): 2.59 ± 0.71 μm (**Fig 4E**); YpsA (+ inducer): 6.36 ± 2.56 μm (**Fig 4F**) | YpsA-GFP (- inducer): 2.99 ± 0.73 μm (**Fig. 4G**); YpsA-GFP (+ inducer): 6.77 ± 3.48 μm (**Fig. 4H**)].

Upon imaging the Δ*yfhS* cells, to our astonishment, we noticed that the average cell length was smaller than WT cells [Δ*yfhS* (- inducer): 1.99 ± 0.61 μm], compare **Figs. 4I** and **4C**. In addition to a smaller cell length, the average cell width of Δ*yfhS* cells also appeared to be smaller when compared to WT [WT (- inducer): 0.78 ± 0.06 μm (**Fig. 4C**); Δ*yfhS* (- inducer): 0.67 ± 0.08 μm (**Fig. 4I**) | n=100]. This observation hints at the possible role for YfhS in cell size regulation either directly or indirectly. Addition of inducer had no effect on the average cell length of cells lacking *yfhS* [Δ*yfhS* (+ inducer): 1.91 ± 0.48 μm; **Fig. 4J**]. When cells containing an inducible copy of either *ypsA* or *ypsA-gfp* in a *yfhS* null background were imaged, they also exhibited smaller cell lengths in the absence of inducer [Δ*yfhS* + YpsA (- inducer): 1.93 ± 0.52 μm (**Fig. 4K**); Δ*yfhS* + YpsA-GFP (- inducer): 2.02 ± 0.55 μm (**Fig. 4M**)], suggesting that that small-cell phenotype is intrinsically linked to the lack of the *yfhS* gene. Intriguingly, overproduction of either YpsA or YpsA-GFP did not result in filamentation in a *yfhS* null background [Δ*yfhS* + YpsA (+ inducer): 1.97 ± 0.56 μm (**Fig. 4L**); Δ*yfhS* + YpsA-GFP (+ inducer): 2.17 ± 0.72 μm (**Fig. 4N)**, indicating that YpsA-mediated cell division inhibition is dependent on YfhS. The ratio of YpsA-GFP-FLAG and SigA in the WT background and *yfhS* null background were similar (0.68 and 0.56 respectively; **Fig. S1C**), suggesting that the elimination of cell division inhibition is not due to defective accumulation of YpsA-GFP. We did observe a 2-fold increase in the ratio of YpsA-FLAG and SigA between WT and Δ*yfhS* background (0.80 and 1.59 respectively; **Fig. S1C**). However, this can be attributed to lower levels of SigA that we have seen reproducibly in this strain background, when the optical density is standardized between the strains tested. Thus, we conclude that the abolition of cell division inhibition in *yfhS* null strain is not due to defective accumulation of YpsA or YpsA-GFP.

We further confirmed that the *yfhS* deletion phenotype is linked specifically to *yfhS*, and not due to any kind of polar effect, by using the complementation strain described earlier. Fluorescence microscopy revealed that the characteristic small-cell phenotype of Δ*yfhS* was no longer observed, even in the absence of inducer, likely due to the leaky expression of *yhfS* in the complementation strain [Δ*yfhS* + *yfhS* (- inducer): 2.23 ± 0.58 μm (**Fig. 4O**)]. When the expression of ectopic *yfhS* was induced by the addition of inducer, the average cell length resembled that of the WT control [Δ*yfhS* + *yfhS* (+ inducer): 2.47 ± 0.65 μm], compare **Figs. 4P** and **4D**. As expected, cells carrying an IPTG-inducible copy of *ypsA* or *ypsA-gfp* in the complementation strain background, in the absence of inducer, appeared similar to the WT control [Δ*yfhS* + *yfhS* + YpsA (- inducer): 2.31 ± 0.76 μm (**Fig. 4Q**); Δ*yfhS* + *yfhS* + YpsA-GFP (- inducer): 2.44 ± 0.80 μm (**Fig. 4S**)]. However, in the presence of inducer filamentation was restored in these two strains [Δ*yfhS* + *yfhS* + YpsA (+ inducer): 5.60 ± 2.47 μm (**Fig. 4R**); Δ*yfhS* + *yfhS* + YpsA-GFP (+ inducer): 7.25 ± 3.64 μm (**Fig. 4T**)], confirming that YpsA-mediated filamentation requires YfhS. The precise reason for this requirement is unclear at this time.

## DISCUSSION

Although many factors involved in facilitating the cell division process have been discovered in *B. subtilis* (Errington & Wu, 2017) and *E. coli* (Du & Lutkenhaus, 2017), our understanding is still incomplete even in these model organisms as evidence of yet to be uncovered factors exists (Rodrigues & Harry, 2012, Bailey *et al.*, 2014). We reported previously that YpsA is such a factor, which appears to play a role in cell division in *B. subtilis* and *S. aureus* (Brzozowski *et al.*, 2019a). The precise mechanism by which YpsA functions remains unclear. The structure of YpsA and another SLOG superfamily member DprA, a single-stranded DNA binding protein, is similar. Thus it is possible YpsA also binds DNA, or nucleotides such as NAD or ADP-ribose as speculated previously (Brzozowski *et al.*, 2019a). We undertook this study to shed light on the possible pathways through which YpsA functions. In this report, we describe our observations of YpsA-mediated lethality on solid medium and utilized that phenomenon as a tool to isolate spontaneous suppressors. Using this technique, we have isolated intragenic suppressors and an extragenic suppressor that abolishes YpsA-mediated toxicity.

We have isolated four intragenic suppressors (E55D, P79L, R111P, G132E) using our screen. Given that E55 and P79 residues are highly conserved among YpsA, perhaps not surprisingly, mutations in those residues render YpsA inactive at least with respect to its function in cell division. It appears that in the E55D mutation, even though it retains the negative charge, shortening of the side chain appears to weaken the ability to form hydrogen bonds with neighboring residues. The mutations in highly conserved P79 and weakly conserved G132 residues create several steric clashes, which explains why the function of YpsA in cell division is affected. The mutation in the solvent exposed residue R111 may result in weakened intramolecular and/or intermolecular interactions. Thus, our screen has identified multiple key residues that are essential for the proper function of YpsA in regard to cell division inhibition.

The extragenic suppressor mutation we isolated introduced a premature stop codon in the *yfhS* open reading frame. YfhS is a relatively small protein (74 amino acids) of unknown function. *yfhS* is upregulated during sporulation through the SigE transcription factor (Yamamoto *et al.*, 1999) and possibly by AbbA (Banse *et al.*, 2008), thus it has been classified as a sporulation gene. In our results we note that Δ*yfhS* cells appear smaller in both width and length compared to our WT control, suggesting that YfhS may have a role in cell size regulation during vegetative growth. Furthermore, we tested and confirmed that YpsA-mediated cell division inhibition is dependent on the presence of full length YfhS.

Given that YfhS is a protein of unknown function, how YfhS and YpsA are linked remains to be determined. During our course of experiments, we noticed that Δ*yfhS* cells grew slower than our WT control (**Fig. S3B**). We have previously shown that YpsA-mediated cell division inhibition is a growth rate-dependent phenomenon (Brzozowski *et al.*, 2019a), thus it is possible that the abolition of cell division inhibition in cells lacking *yfhS* could be attributed to slow growth. However, at this point we cannot rule out the possibility of a direct mechanistic link between YpsA and YfhS, since YpsA plays a role in cell division and YfhS appears to play a role in cell size regulation.

## EXPERIMENTAL PROCEDURES

### Strain construction and general methods

All *B. subtilis* strains utilized during the course of this study are derivatives of the laboratory strain PY79 (Youngman *et al.*, 1984). **Table S1** contains all relevant strain and oligonucleotide information. The construction of strains overexpressing *ypsA, ypsA-gfp, ypsA-flag*, and *ypsA-gfp-flag* have been described previously (Brzozowski *et al.*, 2019a). In order to construct a *B. subtilis* strain containing an inducible copy of *yfhS, yfhS* was PCR amplified from PY79 chromosomal DNA using primer pair oRB59/oRB60. The resulting PCR product was digested with SalI and NheI restriction enzymes and cloned into pDR111 (D. Rudner), also digested with SalI and NheI, to construct plasmid pRB54. The constructed plasmids were then transformed into competent PY79 cells to introduce genes of interest via double crossover homologous recombination into either the native and non-essential *amyE* locus or into a second *amyE* locus (bkdB::Tn917::amyE::cat; Amy Camp).

### Media and culture conditions

Overnight *B. subtilis* cultures were grown at 22 °C in Luria-Bertani (LB) growth medium, and subsequently diluted 1:10 into fresh LB medium. Cultures were grown at 37 °C in a shaking incubator to mid-logarithmic growth phase (OD_600_=0.5), unless otherwise stated. In order to induce the expression of genes under the control of an IPTG-inducible promoter, 250 μM IPTG was added to growing cultures, where required, at mid-logarithmic phase, unless stated otherwise.

### Spot assay

All spot assays were completed on LB agar plates supplemented with 1mM IPTG, where required, to induce the expression of genes under the control of an IPTG-inducible promoter. Required strains were first grown to mid-logarithmic phase (OD_600_=0.5) at 37°C while shaking, and subsequently standardized to an OD_600_=0.1. Following standardization, serial dilutions of each of the strains were spotted onto the appropriate LB plates at a volume of 1 μl. Plates were incubated overnight (approximately 14 hours) at 37 °C. On the following day, plates were observed for growth defects.

### Isolation of spontaneous suppressor mutations

The severe growth defect associated with the strain overproducing YpsA-GFP (GG83) allowed for the isolation of spontaneous suppressor mutations that were able to restore growth similar to the WT control. Suppressor mutations were isolated and determined to be either intragenic or extragenic as indicated in **Fig. S2**. For this purpose, the strain GG83 was plated on LB agar plates containing 1 mM IPTG to induce the expression of *ypsA*-*gfp*, and plates were incubated overnight at 37 °C. After overnight incubation, plates were examined for growth defects associated with the overproduction of YpsA-GFP. PY79 was utilized as a control to ensure that any reduction in growth was specifically due to the overproduction of YpsA-GFP. Single colonies of GG83 that did arise in the presence of inducer (likely containing suppressor mutations) were isolated from the original plate and used to inoculate new LB agar plates that were then grown overnight at 37 °C. Genomic DNA was then isolated from each of the strains containing suppressor mutations using standard phenol-chloroform DNA extractions. Isolated genomic DNA was then used to transform WT PY79 cells, which were then screened for integration of *ypsA-gfp* into the non-essential *amyE* locus. The resulting transformants were then inoculated onto LB agar plates supplemented with 1 mM IPTG to induce the expression of *ypsA-gfp*, and plates were incubated overnight at 37 °C. On the following day plates were screened for growth defects associated with *ypsA-gfp* overexpression. PY79 was used as a control to ensure that any observed growth defect was specifically due to the production of YpsA-GFP. If strains harboring an inducible copy of *ypsA-gfp* isolated during our suppressor screen were now able to grow in the presence of the IPTG inducer, then the suppressor mutations were noted as possibly intragenic (*ypsA*\**-gfp*). If the strains were still unable to grow in the presence of the IPTG inducer, then the mutations were labeled as possibly extragenic - as this indicated that the inducible copy of *ypsA-gfp* within the *amyE* locus did not contain any mutations that were able to restore WT-like growth. All strains determined to contain intragenic suppressor mutations within the IPTG-inducible copy of *ypsA-gfp* at the *amyE* locus were screened via fluorescence microscopy to ensure GFP fluorescence, ruling out some potential mutations within the promoter region, frame-shift mutations, and introduction of premature stop codons. Genomic DNA was isolated from each of the *ypsA*\**-gfp* strains containing intragenic suppressor mutations and was then used as a template for PCR using primer pair oP106/oP24 to amplify the *ypsA-gfp* within the *amyE* locus. The resulting PCR products were sequenced using a 3’ internal GFP sequencing primer (oP212) by GENEWIZ (South Plainfield, NJ). Sequence analysis was completed using ApE Plasmid Editor (v2.0.51) (M. Wayne Davis), and multiple sequence alignments were built using Clustal Omega multiple sequence alignment software (Sievers *et al.*, 2011). All strains characterized as containing extragenic suppressor mutations that restored WT-like growth to strains overproducing YpsA-GFP were subjected to additional screening prior to whole genome sequencing by integrating a new copy of *ypsA-gfp* into the *amyE* locus. The original copy of *ypsA-gfp* was first replaced by a chloramphenicol resistance cassette, and the resulting strain was then transformed with pGG28 (Brzozowski *et al.*, 2019a), to reintroduce a new copy of *ypsA-gfp* into the *amyE* locus. The resulting strains were then used to inoculate LB agar plates supplemented with 1 mM IPTG to verify that they were still able to grow in the presence of inducer unlike the GG83 parental strain. PY79 and GG83 were used as controls on these plates. Plates were incubated overnight at 37 °C and observed for any growth defects associated with YpsA-GFP overproduction on the following day. Genomic DNA was isolated from strains containing extragenic suppressor mutations using the Wizard Genomic DNA Purification Kit (Promega) and sent for whole genome sequencing (Tufts University School of Medicine Genomics Core).

### Bioinformatics and variant detection

Data was analyzed using CLC Genomics Workbench 11 (Qiagen Bioinformatics). First, raw reads were aligned to the PY79 reference sequence (CP00681) using the Map Reads to Reference tool. Output read mappings were then subject to coverage analysis and variant detection. The Basic Variant Detection tool was used to generate a variant track and variant table output in consideration with coverage results. Resultant amino acid changes of variants unique to extragenic suppressors were examined using the Amino Acid Changes tool (version 2.4) using set for genetic code parameter 11: Bacterial, Archaeal, and Plant Plasmid. Subsequently, suppressor mutations were also verified manually.

### Structural analysis

All figures and rotamers were generated using PyMOL (Schrödinger, LLC).

### Growth Curves

PY79, RB314, and RB409 were first grown to mid-logarithmic phase (OD_600_=0.5) in LB broth at 37 °C with shaking and subsequently standardized to an OD_600_=0.1. IPTG was added to the growth medium at a final concentration of 1 mM, where required to induce the expression of genes of interest. Cultures were then grown in LB medium at 37 °C with shaking for a total elapsed time of 6 h. Growth curves were plotted using GraphPad Prism version 8.3.1 (GraphPad Software, La Jolla, California, USA).

### Microscopy

Microscopy was completed by taking 1 ml aliquots of *B. subtilis* cultures and washing with 1X phosphate buffered saline (PBS) through centrifugation. Cells were then resuspended in 100 μl of PBS, and the red membrane stain FM4-64 was added at a final concentration of 1 μg/ml. The sample was prepared for microscopy by spotting 5 μl of the cell suspension onto the glass coverslip of a MatTek glass bottom dish and subsequently covering it with a 1% agarose pad made with sterile water as described previously (Brzozowski *et al.*, 2019b). All imaging was completed at room temperature inside of an environmental chamber using a GE Applied Precision DeltaVision Elite deconvolution fluorescence microscope. Photos were taken using a Photometrics CoolSnap HQ2 camera. All images were acquired by taking 17 Z-stacks at 200 nm intervals. Images were deconvolved though the SoftWorx imaging software provided by the microscope manufacturer.

### Immunoblot Analysis

*B. subtilis* strains were grown overnight at 22 °C in LB growth medium, and then diluted 1:10 into fresh LB the following day. Cultures were grown to an OD_600_=0.5 and subsequently induced with 1 mM IPTG where required to induce the expression of the genes of interest. Cultures were then grown to OD_600_=1.0 and following the induction period, 1 ml aliquots of cultures were centrifuged, and cell lysis was completed by resuspending the cell pellet in a protoplast buffer containing 0.5 M sucrose, 20 mM MgCl_2_, 10 mM KH_2_PO_4_, and 0.1 mg/ml lysozyme. Samples were incubated at 37°C for 30 min and then prepared for SDS-PAGE. Following electrophoresis, samples were transferred onto nitrocellulose membrane and subsequently probed with antibodies against GFP, FLAG (Proteintech Group Inc.), or *B. subtilis* SigA (M. Fujita), which was used as an internal loading control.

## ACKNOWLEDGEMENTS

We thank our lab members for comments on the manuscript. This work was funded by a start-up grant from USF (PE), and the National Institutes of Health grants (R35GM133617; PE) and (R01AI124458; LS).

## AUTHOR CONTRIBUTIONS

The conception and design of the study (RSB, PJE), data acquisition (RSB, JJC, ANA, PJE), analysis and/or interpretation of the data (RSB, BRT, MDS, YC, LNS, PJE), and writing of the manuscript (RSB, MDS, YC, PJE).

## FIGURE LEGENDS

**Table S1.** Strains and oligonucleotides used in this study.

**Figure S1**. Analysis of YpsA accumulation. (**A**) Fluorescence microscopy of strains containing an IPTG-inducible copy of either *ypsA-flag* (RB121) or *ypsA-gfp-flag* (RB125) grown in the absence of inducer (left panels) or in the presence of 250 μM IPTG (right panels). Fluorescence of FM4-64 membrane dye (red) and GFP (green) are shown. Scale bar: 1 μm. (**B**) Spot assays including wild-type cells (PY79) and cells containing an IPTG-inducible copy of either *ypsA* (GG82), *ypsA-gfp* (GG83), *ypsA-flag* (RB121), or *ypsA-gfp-flag* (RB125). Dilutions of standardized cultures were spotted on solid medium without inducer (left panel) or containing 1 mM IPTG (right panel). Corresponding dilution factors are indicated below. (**C**) Anti-FLAG and anti-SigA (loading control) immunoblots of RB121 (YpsA-FLAG), RB412 (Δ*yfhS* + YpsA-FLAG), RB125 (YpsA-GFP-FLAG), and RB413 (Δ*yfhS* + YpsA-GFP-FLAG) cell lysates. The FLAG/SigA ratios corresponding to each lane are shown.

**Figure S2**. Flow chart detailing the methodology used to screen spontaneous suppressor mutations.

**Figure S3**. Analysis of *yfhS* extragenic suppressor mutation. (**A**) Pairwise alignment of the *yfhS* sequence in WT (PY79) and the extragenic suppressor (RBSS6E11). The source of 10-nucleotide duplication is highlighted. (**B**) Growth curves of WT (PY79), Δ*yfhS* (RB314), and *yfhS* complementation strain (RB409) are shown.

