## Supplemental Data for "Suppressors of YpsA-mediated cell division inhibition in *Bacillus subtilis*"

**Table S1.** Strains and oligonucleotides used in this study.

**Figure S1**. Analysis of YpsA accumulation. (**A**) Fluorescence microscopy of strains containing an IPTG-inducible copy of either *ypsA-flag* (RB121) or *ypsA-gfp-flag* (RB125) grown in the absence of inducer (left panels) or in the presence of 250 μM IPTG (right panels). Fluorescence of FM4-64 membrane dye (red) and GFP (green) are shown. Scale bar: 1 μm. (**B**) Spot assays including wild-type cells (PY79) and cells containing an IPTG-inducible copy of either *ypsA* (GG82), *ypsA-gfp* (GG83), *ypsA-flag* (RB121), or *ypsA-gfp-flag* (RB125). Dilutions of standardized cultures were spotted on solid medium without inducer (left panel) or containing 1 mM IPTG (right panel). Corresponding dilution factors are indicated below. (**C**) Anti-FLAG and anti-SigA (loading control) immunoblots of RB121 (YpsA-FLAG), RB412 (Δ*yfhS* + YpsA-FLAG), RB125 (YpsA-GFP-FLAG), and RB413 (Δ*yfhS* + YpsA-GFP-FLAG) cell lysates. The FLAG/SigA ratios corresponding to each lane are shown.

**Figure S2**. Flow chart detailing the methodology used to screen spontaneous suppressor mutations.

**Figure S3**. Analysis of *yfhS* extragenic suppressor mutation. (**A**) Pairwise alignment of the *yfhS* sequence in WT (PY79) and the extragenic suppressor (RBSS6E11). The source of 10-nucleotide duplication is highlighted. (**B**) Growth curves of WT (PY79), Δ*yfhS* (RB314), and *yfhS* complementation strain (RB409) are shown.

**Table S1 Strains and oligonucleotides used in this study**

Strains used in this study

| **Strain** | **Genotype** | **Reference** |
|---|---|---|
| PY79 | Wild type | Youngman *et al.* (1984) |
| GG82 | *amyE::P*$_{hyperspank}$*-ypsA spec* | Brzozowski *et al.* (2019) |
| GG83 | *amyE::P*$_{hyperspank}$*-ypsA-gfp spec* | Brzozowski *et al*. (2019) |
| RB121 | *amyE::P*$_{hyperspank}$*-ypsA-flag spec* | Brzozowski *et al*. (2019) |
| RB125 | *amyE::P*$_{hyperspank}$*-ypsA-gfp-flag spec* | Brzozowski *et al*. (2019) |
| RB300 | *amyE::P*$_{hyperspank}$*-ypsA*$^{G132E}$*-gfp spec* | This study |
| RB301 | *amyE::P*$_{hyperspank}$*-ypsA*$^{P79L}$*-gfp spec* | This study |
| RB327 | *amyE::P*$_{hyperspank}$*-ypsA*$^{E55D}$*-gfp spec* | This study |
| RB328 | *amyE::P*$_{hyperspank}$*-ypsA*$^{R111P}$*-gfp spec* | This study |
| RB314 | Δ*yfhS::erm* | Derived from BKE08640 (BGSC*) |
| RB288 | Δ*yfhS::erm; amyE::P*$_{hyperspank}$*-ypsA spec* | This study |
| RB289 | Δ*yfhS::erm; amyE::P*$_{hyperspank}$*-ypsA-gfp spec* | This study |
| RB409 | Δ*yfhS::erm; bkdB::Tn917::amyE::cat::P*$_{hyperspank}$*-yfhS spec* | This study |
| RB410 | Δ*yfhS::erm; bkdB::Tn917::amyE::cat::P*$_{hyperspank}$*-yfhS spec; amyE::P*$_{hyperspank}$*-ypsA spec::cat* | This study |
| RB411 | Δ*yfhS::erm; bkdB::Tn917::amyE::cat::P*$_{hyperspank}$*-yfhS spec; amyE::P*$_{hyperspank}$*-ypsA-gfp spec::cat* | This study |
| RB412 | Δ*yfhS::erm; amyE::P*$_{hyperspank}$*-ypsA-flag spec* | This study |
| RB413 | Δ*yfhS::erm; amyE::P*$_{hyperspank}$*-ypsA-gfp-flag spec* | This study |
| RBSS6E11 | *amyE::P*$_{hyperspank}$*-ypsA-gfp*** spec* | This study |

* Bacillus Genetic Stock Center

** Strain carries suppressor mutation

Oligonucleotides used in this study

| **Primer** | **Sequence (5' to 3')** |
|---|---|
| oP24 | GCCGCATGC*TTA* TTTGTATAGTTCATCCATGCC |
| oP106 | AAAGTCGAC*ACATAAGGAGGAACTACT* ATGAAAGTATTGGCAATAACGGGCTATAAACCG |
| oP212 | GGGTAAGTTTTCCGTATGTTGCATCACCTTCACCCTCTCC |
| oRB59 | AATAAGTCGAC*ACATAAGGAGGAACTACT* ATGTATGTCGGACGTGATATGAGCGAA |
| oRB60 | AATAAGCTAGC*TTA* ATCGTAAGAGACGCGCGTGCCGTGGCT |

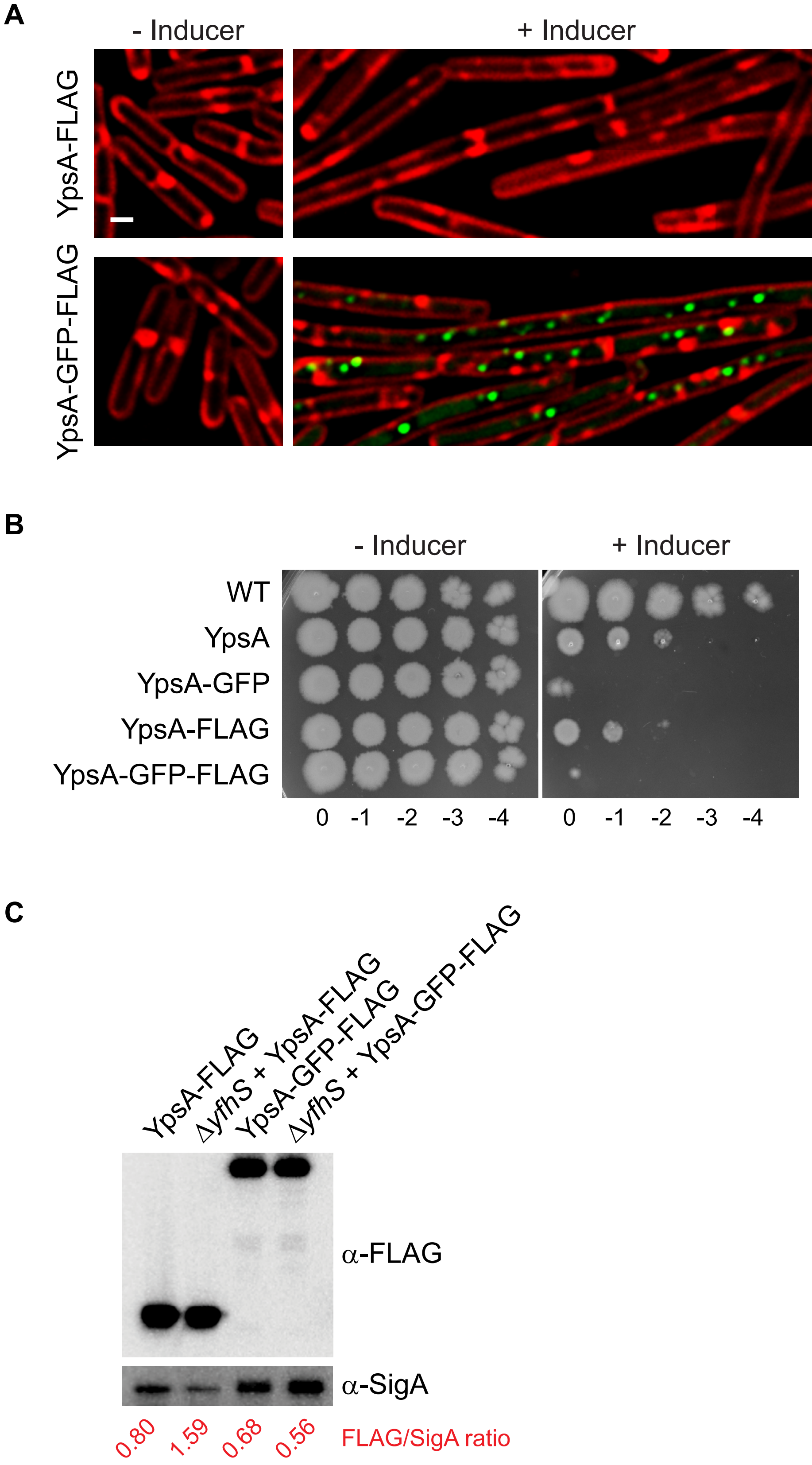
A
- Inducer
+ Inducer
YpsA-FLAG
YpsA-GFP-FLAG
B
- Inducer
+ Inducer
WT
YpsA
YpsA-GFP
YpsA-FLAG
YpsA-GFP-FLAG
0 -1 -2 -3 -4
0 -1 -2 -3 -4
C
YpsA-FLAG
Δ*yfhS* + YpsA-FLAG
YpsA-GFP-FLAG
Δ*yfhS* + YpsA-GFP-FLAG
α-FLAG
α-SigA
0.80
1.59
0.68
0.56
FLAG/SigA ratio

Figure S1

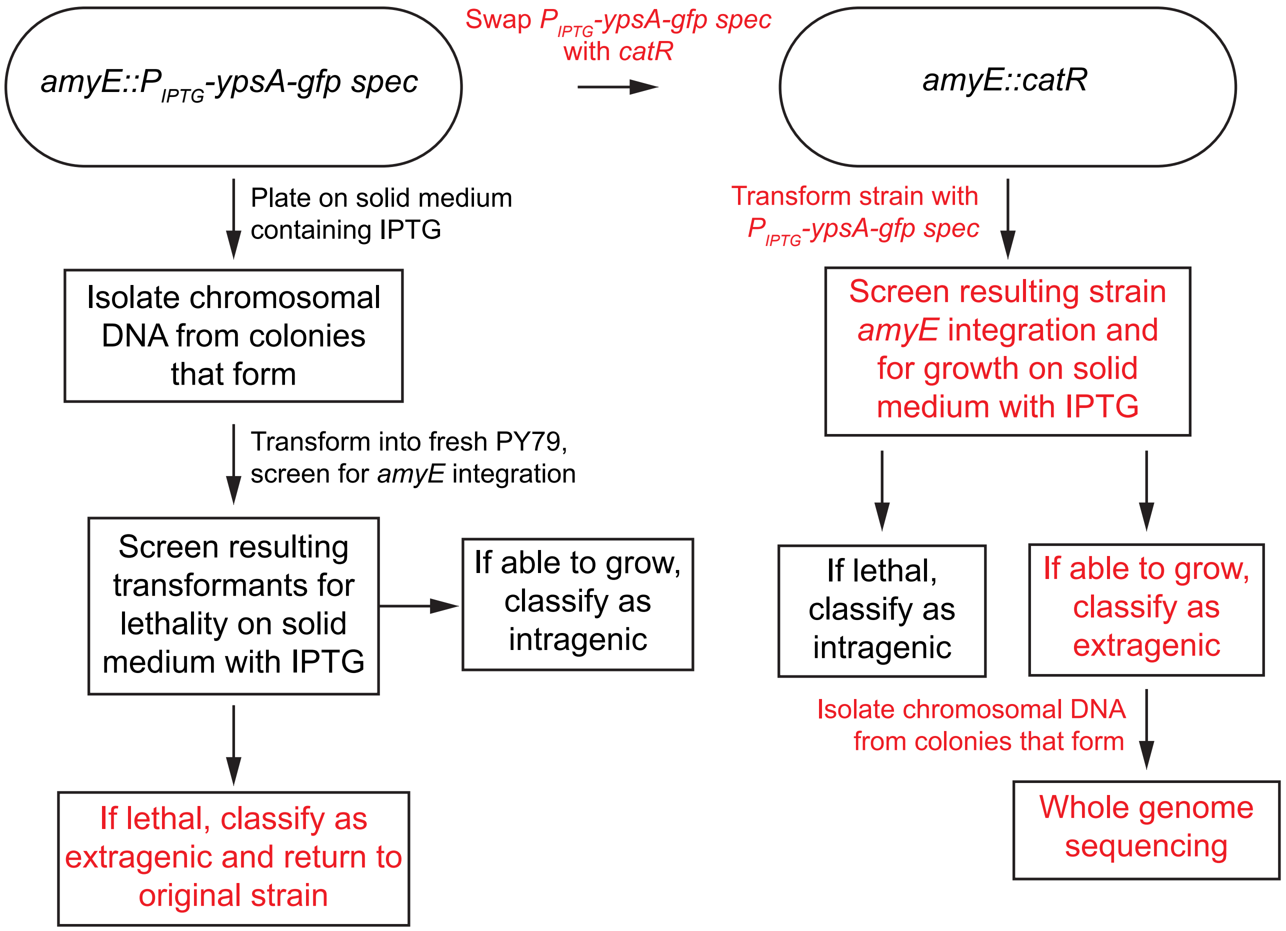
amyE::P$_{IPTG}$-ypsA-gfp spec
Swap P$_{IPTG}$-ypsA-gfp spec with catR
amyE::catR
Plate on solid medium containing IPTG
Isolate chromosomal DNA from colonies that form
Transform into fresh PY79, screen for amyE integration
Screen resulting transformants for lethality on solid medium with IPTG
If able to grow, classify as intragenic
If lethal, classify as extragenic and return to original strain
Transform strain with P$_{IPTG}$-ypsA-gfp spec
Screen resulting strain amyE integration and for growth on solid medium with IPTG
If lethal, classify as intragenic
If able to grow, classify as extragenic
Isolate chromosomal DNA from colonies that form
Whole genome sequencing

Figure S2

**A**

```
WT          ATGTATGTCGGACGTGATATGAGCGAATTGAACATGGTTTCCAAAAAAGATTGGAAGAAC   60
RBSS6E11    ATGTATGTCGGACGTGATATGAGCGAATTGAACATGGTTTCCAAAAAAGATTGGAAGAAC   60
            ************************************************************

WT          AGTGAACTCGCTTATTTTCATCATGCCTTTCAGCAAATTATGCCTTATTTGAACGAAGAA   120
RBSS6E11    AGTGAACTCGCTTATTTTCATCATGCCTTTCAGCAAATTATGCCTTATTTGAACGAAGAA   120
            ************************************************************

WT          GGCCAATCAAAATACCGGGAATTAACGCAAGAAATTGAAGCGCGCGGCGGAATGAAGCGC   180
RBSS6E11    GGCCAATCAAAATACCGGGAATTAACGCAAGAAATTGAAGCGCGCGGCGGAATGAAGCGC   180
            ************************************************************

WT          AATGAAG----------CGGACTACAGCCACGGCACGCGCGTCTCTTACGATTAA        225
RBSS6E11    AATGAAGCGCAATGAAGCGGACTACAGCCACGGCACGCGCGTCTCTTACGATTAA        235
            *******          **************************************
```

**B**

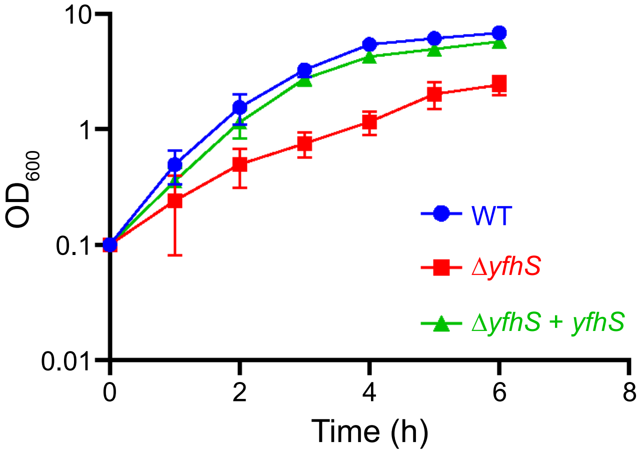


Figure S3
